# DIPLER: rapid k-mer-based locus identification

**DOI:** 10.64898/2026.10.09.757889

**Authors:** Kaur Robin Kivirüüt, Lauris Kaplinski, Andres Salumets, Tarmo Puurand

**Affiliations:** Genetics and Personalized Medicine Clinic, University of Tartu, Tartu, Estonia; Institute of Molecular and Cell Biology, University of Tartu, Tartu, Estonia; Department of Physiology and Pharmacology, Karolinska Institutet, Stockholm, Sweden; Department of Obstetrics and Gynaecology, Institute of Clinical Medicine, University of Tartu, Tartu, Estonia; Celvia CC, Competence Centre on Health Technologies, Tartu, Estonia

**Keywords:** k-mer, locus discovery

## Abstract

Here, we present Dipler, a two-step strategy that combines the identification of group-specific k-mers from whole-genome sequencing data with de novo assembly of PacBio reads enriched for those k-mers. Using a Gg/gg presence/absence-like model in *Juglans hindsii*, we recovered the structurally distinct “G” allele from PacBio reads at coverage as low as 9×. Using the exceptional GG genotype of *J. regia* “Sharkey” as a complementary diploid model, we further show that two divergent “G” alleles of the same locus can be resolved and assembled separately from 13× coverage. Dipler therefore provides a rapid, reference-independent framework for identifying structurally complex loci, with potential applicability across eukaryotic species.

## INTRODUCTION

Methods for identifying causal loci have evolved alongside approaches for detecting and characterising genetic variation. Early studies relied on highly informative markers, such as STRs, together with linkage maps^1^, making locus discovery laborious and slow compared with current methods. The emergence of reference genomes transformed this process by providing a much richer representation of genomic context than sparse marker panels could offer. In humans, this transition occurred roughly two decades ago, and reference-based analysis subsequently became the dominant framework^2^. However, with current sequencing technologies, that framework is no longer universally optimal, because whole-genome sequencing is now far more accessible and can capture forms of variation that are incompletely represented in reference panels^3^.

Whole-genome sequencing enables both GWAS-based and k-mer-based locus discovery strategies. Although GWAS has been extensively validated in association studies, k-mer-based approaches remain comparatively underdeveloped. The availability of telomere-to-telomere assemblies has nevertheless improved the interpretability of k-mer-based analyses by facilitating the genomic mapping of informative sequences^4^. In addition, biologically defined groups often contain internal heterogeneity^4^, creating a setting in which discriminating k-mers can be detected using non-parametric tests such as the Mann– Whitney test.

Here, we extend the use of k-mers beyond the Y-mer framework^4^ and evaluate Dipler using two complementary models of locus recovery. First, in *J. hindsii*, we use the Gg/gg contrast to identify k-mers associated with flowering order and use these k-mers to select PacBio reads for de novo assembly of the structurally distinct “G” allele. Because much of the “G”-specific sequence is absent from the homologous “g” allele, this represents a presence/absence-like model in which the target sequence is present on only one homolog in Gg individuals. We then use the flowering-order-discriminating k-mers identified in *J. hindsii* to classify *J. regia* samples and identify k-mers characteristic of the full “G” locus. Finally, we use the exceptional GG genotype of *J. regia* “Sharkey” as a complementary diploid model to test whether Dipler can resolve and assemble two divergent “G” alleles of the same locus separately. Together, these two models allow us to evaluate locus recovery both when the target sequence is largely absent from one homolog and when two divergent alleles of the same diploid locus must be distinguished and assembled.

## RESULTS

### Phenotype selection

Our aim was to develop a k-mer-based method for identifying phenotype-determining loci in genomes, including regions that are generally considered difficult to analyse. As a proof of concept, we used short-read WGS samples from *Juglans hindsii*, for which flowering-order phenotypes have been described and the underlying locus has been identified^5^. In *J. hindsii*, flowering order is determined by two alleles on chromosome 11: the “g” allele confers male-first flowering, whereas the dominant “G” allele confers female-first flowering^5^.

The “G” and “g” loci differ in the copy number of an approximately 1-kbp repeat: the “g” allele contains a single copy, whereas the “G” allele contains 8–12 copies^5^. This architecture makes the locus structurally complex and therefore well suited for evaluating the method as a proof of concept.

### Increasing group size improves k-mer specificity

To determine the optimal number of samples for locus identification, we tested group-size combinations of 7+7, 8+8, 9+9, 10+10, and 11+11 (Tables 1 and S1). As hypothesised, smaller groups contained larger numbers of shared k-mers that were absent from the opposite group. This pattern was confirmed by our selection criterion: the 7+7 combination yielded 19,089 k-mers specific to the “G” group, whereas the 11+11 combination yielded 7,131 (Table S2). We then examined the correspondence of the identified k-mers both in the reads used to construct the reference genome and in the reference genome itself at the locus identified in another project (Table S3). With only a few exceptions, the “G”-specific k-mers were present in the reads used for reference construction. Moreover, in the reference genome, 60.2% of the k-mers identified in the 7+7 group and 99.7% of those identified in the 11+11 group mapped to the correct chromosome (Table 1).

**Table 1.**
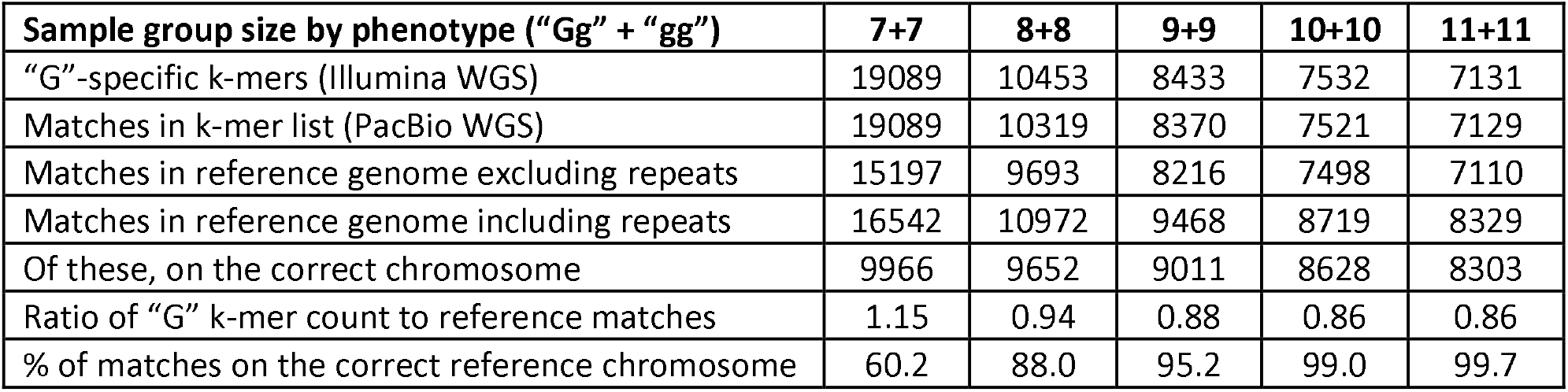
k-mer statistics by phenotype-group size. In the 11+11 group, 99.7% of “G”-group-specific k-mers map to the correct chromosome in the reference sequence (Table S3).

In contrast, *de novo* contigs generated using k-mers identified in randomly assigned sample groups (154 and 160) did not contain most of the selected k-mers (Table S4). The similar magnitude of these results indicates the presence of randomly shared k-mers among the 11+11 samples.

### Reference-free assembly of the J. hindsii Gg/gg presence/absence-like model

The search for group-specific k-mers indicated that 10+10 was an effective sample-size combination. Using the grep function in SeqKit^6^, we identified reads containing the 7,532 “G”-specific k-mers from the 10+10 analysis; in total, 3,817 reads were recovered (Table S5). To determine the minimum PacBio coverage at which the target “G” locus could be recovered, the *J. hindsii* dataset was downsampled to generate read sets with progressively increasing coverage. Each read was then assigned to a coverage group according to both its position in the sequencing dataset and the order of the sequencing run (Table S5). In the Gg/gg presence/absence-like model, the target “G” locus was recovered from the de novo assembly at 9× coverage when reads were added according to sequencing-run order. When the same dataset was downsampled by adding reads according to their order in the SRA file, the target locus was first recovered at 13× coverage (Table S6). Thus, the difference between the 9× and 13× values reflects the different read sets generated by the two downsampling procedures at the same nominal coverage. These coverage values are empirical observations from the respective downsampled read sets and should not be interpreted as fixed minimum coverage thresholds for locus recovery. As coverage increased, Hifiasm^7-9^ produced more contigs; however, with the exception of the target contig, the representation of group-specific k-mers remained low (Table S6).

### Reference-free assembly of the J. regia “GG” allele

Because the sequencing dataset did not include information on “Gg” and “gg” allele status in *J. regia*, we inferred these classes using the 10+10 *J. hindsii* k-mers. Presence of the “G” allele was determined based on two clusters of samples with k-mer values >0, with the larger group classified as “Gg” and the smaller group as “gg” (Tables S7 and S8). Using a group size of 11+11, we identified 7,674 k-mers characteristic of the “G” group (Table S9), most of which mapped to chromosome 11 of the *J. regia* reference genome (Table S10). For the PacBio analysis, we selected J. regia “Sharkey” because this cultivar carries the exceptional “GG” genotype, with two divergent “G” alleles^10,11^. This provided a complementary diploid locus-recovery model for testing whether Dipler could distinguish and assemble two divergent alleles of the same locus. Hifiasm recovered a single “G” allele at 6×–10× coverage. At 11× and 12×, two similar assemblies representing the same allele were recovered, whereas the two divergent “G” alleles were first resolved and assembled separately at 13× coverage and remained separately resolved at higher coverages (Table S11). Thus, 13× represents the empirical coverage at which both divergent alleles were resolved in this dataset and should not be interpreted as a general minimum coverage requirement for Dipler.

### Alignment of J. regia “GG” de novo assemblies

To assess the correctness of the de novo assemblies across different sequencing coverages, we performed a ClustalW^12^ multiple alignment of contigs assembled from 6×–14× dilutions of the J. regia “Sharkey” cultivar (Table S11), in which “G”-group-specific k-mers had been identified. The alignment showed that at 6×–12× coverage, a contig was assembled for only one allele, although both HAP1 and HAP2 were present at 11× and 12×. From 13× coverage onward, the second “G” allele was also assembled separately, and in the ClustalW output these assemblies carried the same sequence differences relative to the first “G” allele (Figure 1).

**Figure 1.**
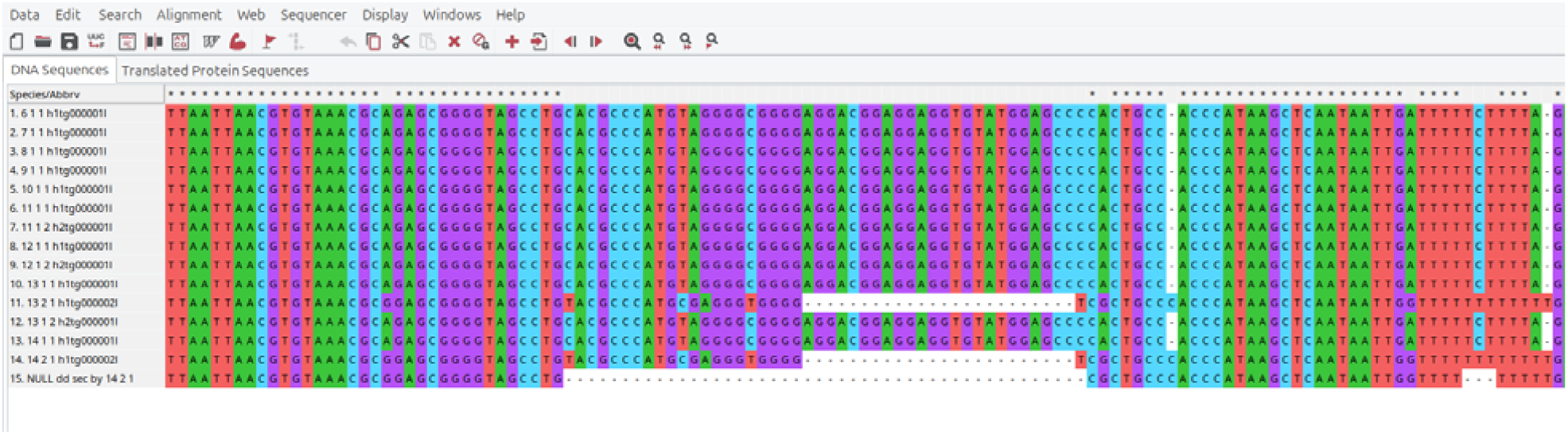
Multiple sequence alignment of de novo-assembled “G” alleles from downsampled read sets. Screenshot from MEGA12^13^ showing a representative region with clear sequence differences between the two “G” alleles. Sequence labels indicate sequencing coverage (6×–14×), haplotype number, contig number (Table S12), and the contig identifier from the Hifiasm output. Sequence differences between the two alleles extend throughout the alignment. The bottom row was added manually and is not part of the MEGA12 alignment output; it visualizes the positions covered by “G”-locus-specific k-mers in the 14_2_1 assembly. These k-mers span positions 12,480–36,943, defining an approximately 24.5-kbp region. Within this region, 7,712 positions are not covered by the selected “G”-locus-specific k-mers because the corresponding k-mers also occur elsewhere in the genome and were therefore excluded during k-mer selection (Supplement).

### Dipler workflow

Whereas the preceding analyses focused on determining the minimum number of group members and the required PacBio sequencing depth, the purpose of the Dipler workflow is to provide a practical and easy-to-use tool. Successful use of the first half of the workflow requires defining groups of samples sequenced at standard coverage and applying a series of GenomeTester4 functions^14^. In the second half, determining the sequence of a complex locus requires at least one PacBio sequencing run, and only after GenomeTester4-based filtering does it become clear from which group the PacBio sample should be selected. Once the relevant subset of reads has been identified, a *de novo* assembler is applied; in this workflow, we used Hifiasm. The stages of the workflow and the required resources are summarised in Figure 2.

**Figure 2.**
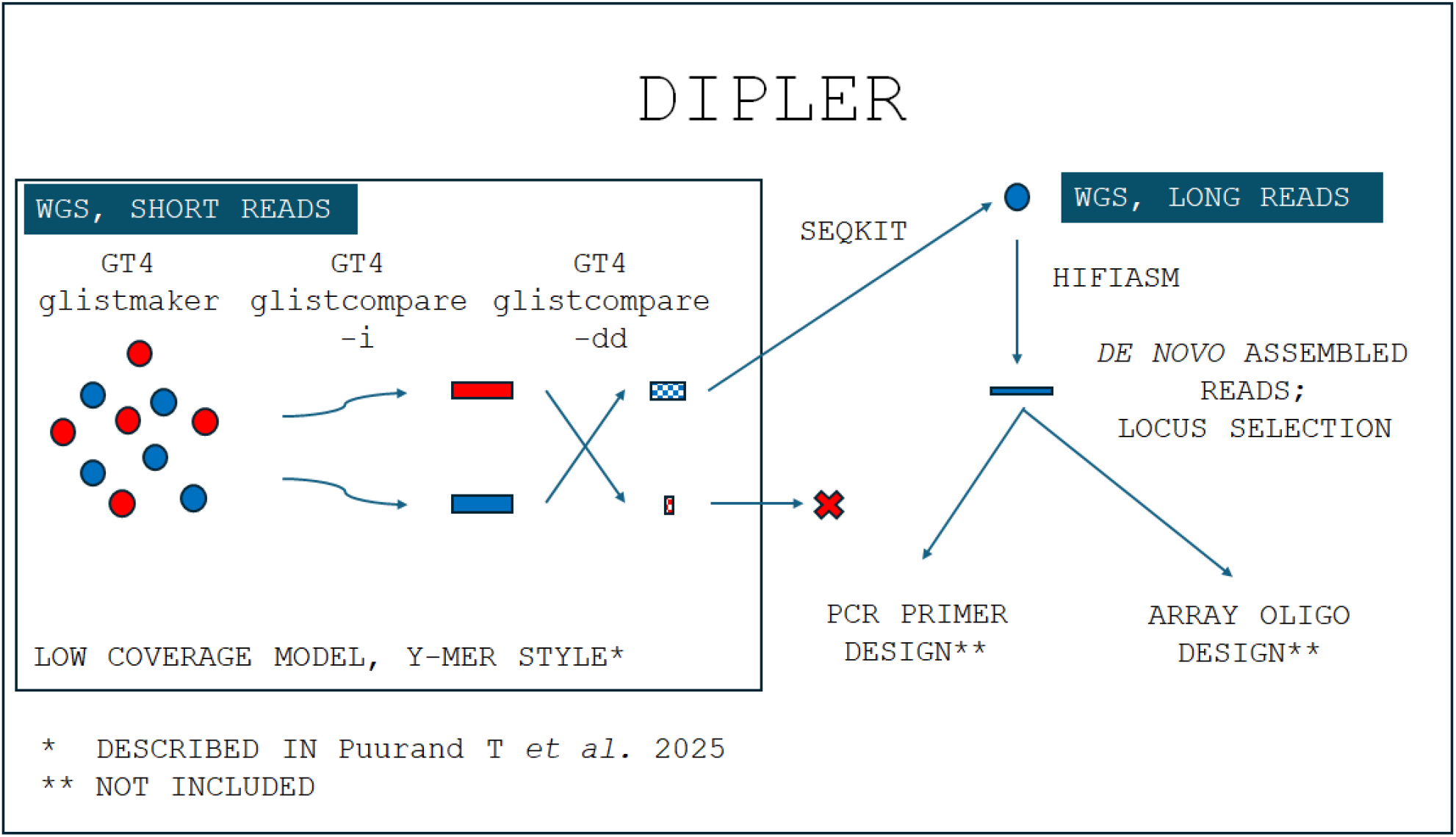
Dipler workflow. GenomeTester4^14^ functions are central to k-mer selection. Using group-specific informative k-mers, the locus of interest is assembled de novo from selected PacBio reads, enabling downstream diagnostic applications.

## DISCUSSION

Our results show that a binary trait with a strong genetic signal can be resolved without a reference-guided discovery workflow. By removing randomly shared k-mers arising from insufficient phenotype representation and filtering out reads carrying only a small number of phenotype-specific k-mers, we recovered the full locus directly from sequencing data. The resulting workflow is rapid and practical.

A key requirement was an adequate group size. We found that 11 samples per group provided a robust balance between specificity and interpretability. Smaller groups yielded more apparently group-specific k-mers, but these additional sequences made locus identification less reliable.

The two models represent biologically distinct locus-recovery challenges and consequently differ in their sequencing-depth requirements. In the *J. hindsii* Gg/gg presence/absence-like model, much of the “G”-specific sequence is absent from the homologous “g” allele, allowing the structurally distinct “G” allele to be recovered at coverage as low as 9×. In contrast, the *J. regia* “Sharkey” GG model requires two divergent “G” alleles of the same diploid locus to be distinguished and assembled separately. This more demanding locus-recovery task required 13× PacBio coverage. These coverage levels are empirical results obtained from the two models examined here and should not be interpreted as universal minimum coverage thresholds for Dipler. Thus, the sequencing depth required for Dipler depends on the underlying locus architecture and on whether the objective is to recover a structurally distinct sequence or to resolve two divergent alleles of the same locus.

Conceptually, Dipler is related to the Y-mer workflow but extends it by incorporating *de novo* assembly of the locus identified by discriminating k-mers. Not all binary-trait loci will be equally tractable: for example, the human Y chromosome is structurally far more complex than a 24.5kbp region. Nevertheless, the framework should be extendable to settings with more than two alleles. In contrast to Y-mer, where repetitive-sequence abundance was a primary focus, Dipler retains only k-mers absent from the comparison group.

Together, these results establish Dipler as a rapid approach for identifying structurally complex loci underlying binary traits. Extending this strategy to multifactorial traits will be an important next step.

## METHODS

### k-mer processing: GListMaker, GListQuery and GListCompare

The GenomeTester4 software package was used for k-mer processing. GenomeTester4 comprises three programs: (1) GListMaker, which creates a sorted list of k-mers from FASTA and FASTQ files; (2) GListCompare, which compares two k-mer lists and identifies their intersection, union, or differences; and (3) GListQuery, which queries k-mer frequencies within a list.

### Identification of group-specific k-mers

Samples were divided into two groups according to flowering order. First, a list of all k-mers together with their frequencies was generated for each individual. Based on the earlier Y-mer study, a k-mer length of 25 was used. Low-frequency k-mers were removed using the minimum at the inflection point of the k-mer-frequency distribution as the threshold. Next, a list of k-mers shared by all members of each group was generated, after which all k-mers shared by both groups were removed. Frequencies of the remaining k-mers were then normalised in individual samples according to sequencing depth. Finally, for each group, characteristic k-mers were identified as those occurring at a higher frequency in individuals of that group than in individuals of the other group, using the Mann–Whitney test.

### de novo assembly workflow

1. Reads containing 10+10 “G”-specific k-mers were identified using SeqKit.

~~~
seqkit grep -f NULL_dd_seq.txt -s SRR29405709_subreads.fastq.gz >
SRR29405709_NULL_dd_seq.fastq
~~~

2. Using an in-house script, all PacBio reads were assigned to sequencing-coverage groups by summing read lengths according to both SRA row number and PacBio run order.
3. Reads containing 10+10 “G”-specific k-mers were assembled *de novo* in onefold coverage increments.

~~~
hifiasm -o katvus/katvus -t4 -f0 katvus_NULL_dd_seq.fastq 2 > katvus/katvus.loq
~~~

4. Using an in-house script, a *.fa file was generated from the *.gfa file; a GT4 list was then indexed, and the number of 10+10 “G”-specific k-mers present in the assembled sequences was determined.

~~~
glistmaker hifiasm.hap1.fasta -w 25 --index -o hifiasm.hap1
glistquery hifiasm.hap1_25.index -f NULL_dd_seq.txt --locations |perl chromosomes_distribution.pl |sort -n -r > hifiasm.hap1.txt
~~~

chromosomes_distribution.pl code

~~~
$jrk = 0;
while(<>){
     chomp;
     @tmp = split(/\t/);
     $jrk++ if $tmp[2] == 0;
     $tabel[$jrk][0] = $tmp[0] if $tmp[2] == 0;
     $tabel[$jrk][$tmp[1]]++;
     $sum{$tmp[1]}++ if ($tmp[0] == 0 && $tmp[3] ne ““);
}
foreach $chr (keys %sum){
     print”$chr\t$sum{$chr}\n”;
}
~~~

5. The output file contains the two columns shown in Table S6: contig number and k-mer count.
6. The correct contig was identified as the one containing the majority of group-specific k-mers.

## Supporting information

alignment file

## DECLARATIONS

### Availability of Data and Materials

*Juglans hindsii*https://www.ebi.ac.uk/ena/browser/view/PRJNA1109986https://trace.ncbi.nlm.nih.gov/Traces/?run=SRR29405709 samples: https://www.ebi.ac.uk/ena/browser/view/PRJNA1109986; https://trace.ncbi.nlm.nih.gov/Traces/?run=SRR29405709

*Juglans hindsii*https://www.ncbi.nlm.nih.gov/datasets/genome/GCA_041380745.1/ “Rawlins”: https://www.ncbi.nlm.nih.gov/datasets/genome/GCA_041380745.1/

*Juglans regia*https://www.ebi.ac.uk/ena/browser/view/PRJNA356989 samples: https://www.ebi.ac.uk/ena/browser/view/PRJNA356989

*Juglans regia*https://www.ncbi.nlm.nih.gov/datasets/genome/GCA_041380835.1/https://www.ncbi.nlm.nih.gov/bioproject/PRJNA1128727/https://trace.ncbi.nlm.nih.gov/Traces/?run=SRR29405708 “Sharkey”:

https://www.ncbi.nlm.nih.gov/datasets/genome/GCA_041380835.1/;

https://www.ncbi.nlm.nih.gov/bioproject/PRJNA1128727/;

https://trace.ncbi.nlm.nih.gov/Traces/?run=SRR29405708

## Authors’ Contributions

T.P. initiated the project. T.P., L.K., and A.S. developed the project design and final concept. T.P. and K.R.K. managed and processed the data. T.P., K.R.K., and L.K. wrote the first draft of the manuscript. K.R.K. wrote the Dipler pipeline. All authors contributed to editing the manuscript. A.S. coordinated and funded the study.

## Acknowledgements

We thank Märt Möls for statistical support and Tõnis Örd for suggestions on the manuscript. Data analysis was performed in part using the facilities of the High-Performance Computing Center at the University of Tartu.

